# Adhesion strength between cells regulate non-monotonic growth by a biomechanical feedback mechanism

**DOI:** 10.1101/2021.11.18.469073

**Authors:** Abdul N Malmi-Kakkada, Sumit Sinha, Xin Li, D. Thirumalai

**Affiliations:** Department of Chemistry and Physics, Augusta University, Augusta, GA 30912, USA; Department of Physics, University of Texas at Austin, Austin, TX 78712, USA; Department of Chemistry, University of Texas at Austin, Austin, TX 78712, USA

## Abstract

We probe the interplay between intercellular interactions and pressure fluctuations associated with single cells in regulating cell proliferation using simulations of a minimal model for three-dimensional multicellular spheroid (MCS) growth. The emergent spatial variations in the cell division rate, that depends on the location of the cells within the MCS, is regulated by intercellular adhesion strength (*f*^*ad*^). This in turn results in non-monotonic proliferation of cells in the MCS with varying adhesion strength, which accords well with experimental results. A biomechanical feedback mechanism coupling the *f*^*ad*^ and cell-dependent pressure fluctuations relative to a threshold value (*p*_*c*_) determines the onset of a dormant phase, and explains the non-monotonic proliferation response. Increasing *f*^*ad*^ from low values enhances cell proliferation because pressure on individual cells is smaller compared to *p*_*c*_. In contrast, at high *f*^*ad*^, cells readily become dormant and cannot rearrange effectively, leading to arrested cell proliferation. Our work, which shows that proliferation is regulated by pressure-adhesion feedback loop, may be a general feature of tumor growth.

## Introduction

Regulation of cell division is necessary for robust tissue growth and morphogenesis, with rapid cell proliferation being an integral phenotype characterizing early development and tumor growth.^1,2^ Mechanical forces influence cell division and spatial organization of cells through mechano-sensitive processes. ^3–7^ For instance, spatial constraints due to cell spatial packing limits cell proliferation.^8–13^ The spatiotemporal arrangement of cells in response to local force (or stress) fluctuations, arising from intercellular interactions and how it controls cell proliferation, is unclear. Studies suggest^11,14–17^ that the cross-talk between cell mechanical interactions and proliferation is mediated by cell-cell adhesion receptors.

An important mediator of inter cellular adhesive force is cadherin, which plays a critical role in morphogenesis, tissue healing, and tumor growth. ^18,19^ The importance of cadherins is documented through their fundamental importance in maintaining multicellular structure during morphogenesis. ^20,21^ Amongst the family of cell adhesion molecules, E-cadherin is the most abundant, and is expressed in most metazoan cohesive tissues.^22,23^ Mechanical coupling between the cortical cytoskeleton and cell membrane involves the cadherin cytoplasmic domain, with forces exerted across the cell-cell contacts being transmitted between cadherin extracellular domain and the cellular cytoskeletal machinery through the cadherin/catenin complex.^24,25^ Thus, to understand how adhesive forces control the spatial organization of cells within 3D multicellular spheroid (MCS) and determine proliferation, we study the impact of varying intercellular adhesion strength on cell proliferation arising from a pressure dependent biomechanical feedback using a computational model.

We simulate an agent based minimal model (see^26^ for a review) for 3D MCS, where individual cells grow, move in response to intercellular forces and local pressure fluctuations, undergo division, as well as dynamically enter into a dormant phase. We show that variation in cell-cell adhesion strength, *f*^*ad*^, results in non-monotonic growth of the MCS into the surrounding medium. We discover that the pressure experienced by individual cells due to mechanical interactions with its neighbors quantitatively explains the non-monotonic proliferation pattern. The *f*^*ad*^ parameter is a proxy for cell-cell adhesion strength, which could vary due to either changes in the cadherin expression level, cadherin clustering,^27^ increasing time of contact between cells,^27^ or “mechanical polarization”, where cells reorganize their adhesive and cytoskeletal machinery to suppress actin density along cell-cell contact interfaces.^28,29^ By building on the integral feedback mechanism that couples cell dormancy and local pressure,^6,7,11^ we show that the pressure exerted by cells on one another explains the experimentally observed non-monotonic cell collective growth as *f*^*ad*^ is varied from a low to high value. The proposed mechanism could be a general attribute for the growth of MCS.

## Methods

We simulate the collective movement of cells using a minimal model of an evolving tumor embedded in a matrix using an agent-based 3D model.^30–33^ The cells, embedded in a highly viscous material mimicking the extracellular material, are represented as soft, spherical objects that interact via direct elastic (repulsive) and adhesive (attractive) forces.

Cell-to-cell and cell-to-matrix damping account for the effects of friction due to other cells, and the extracellular matrix, such as collagen matrix, respectively. The model explicitly accounts for apoptosis, cell growth and division. We also implicitly take into account the ability of individual cells to transition from growth to dormant phase depending on the local pressure. Thus, the collective motion of cells is determined by systematic cell-cell forces and the dynamics due to stochastic cell birth and apoptosis subject to a free boundary condition. The details of the models are presented in the Supplementary Information (SI) which is primarily based on our previous work.^34–37^

### Assessment of the values of *f*^*ad*^

We estimated the *f*^*ad*^ from the typical strength of cell-cell attractive interactions reported in previous experimental studies in order to assess if they are in the reasonable range. Early experiments showed that the interaction strength between cell adhesion proteoglycans is ∼ 2 × 10^−5^*µN/µm*^2^.^38^ Recently, single cell force spectroscopy has been used to measure the typical forces required to rupture E-cadherin mediated bonds between cells.^39^

A typical force-distance curve (FDC), a plot of 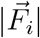 versus 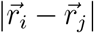 from Eq. S.1 and the elastic force, is shown in Fig. 1**(A)**. The plot shows that, for typical cell sizes (∼ 5*µm*), the minimum force is ≈ 2 × 10^−4^*µN*, which is fairly close to the values ≈ 1.5 × 10^−4^*µN* in Fig. 1**(B)** and 4 × 10^−4^*µN* reported elsewhere.^38^ The inset in Fig. 1**(B)** shows the work (the area under the FDC) required to overcome the inter cellular adhesion force. The value of work done ≈ 0.78 × 10^−16^Nm for primordial germ cells (PGCs in *Xenopus laevis* embryo) is comparable to experiment, and accord well with the energy 1.25 × 10^−16^Nm required to rupture the adhesive forces between two cells used in the simulations. Note that we did not adjust any parameters to obtain reasonable agreement.

**Figure 1:**
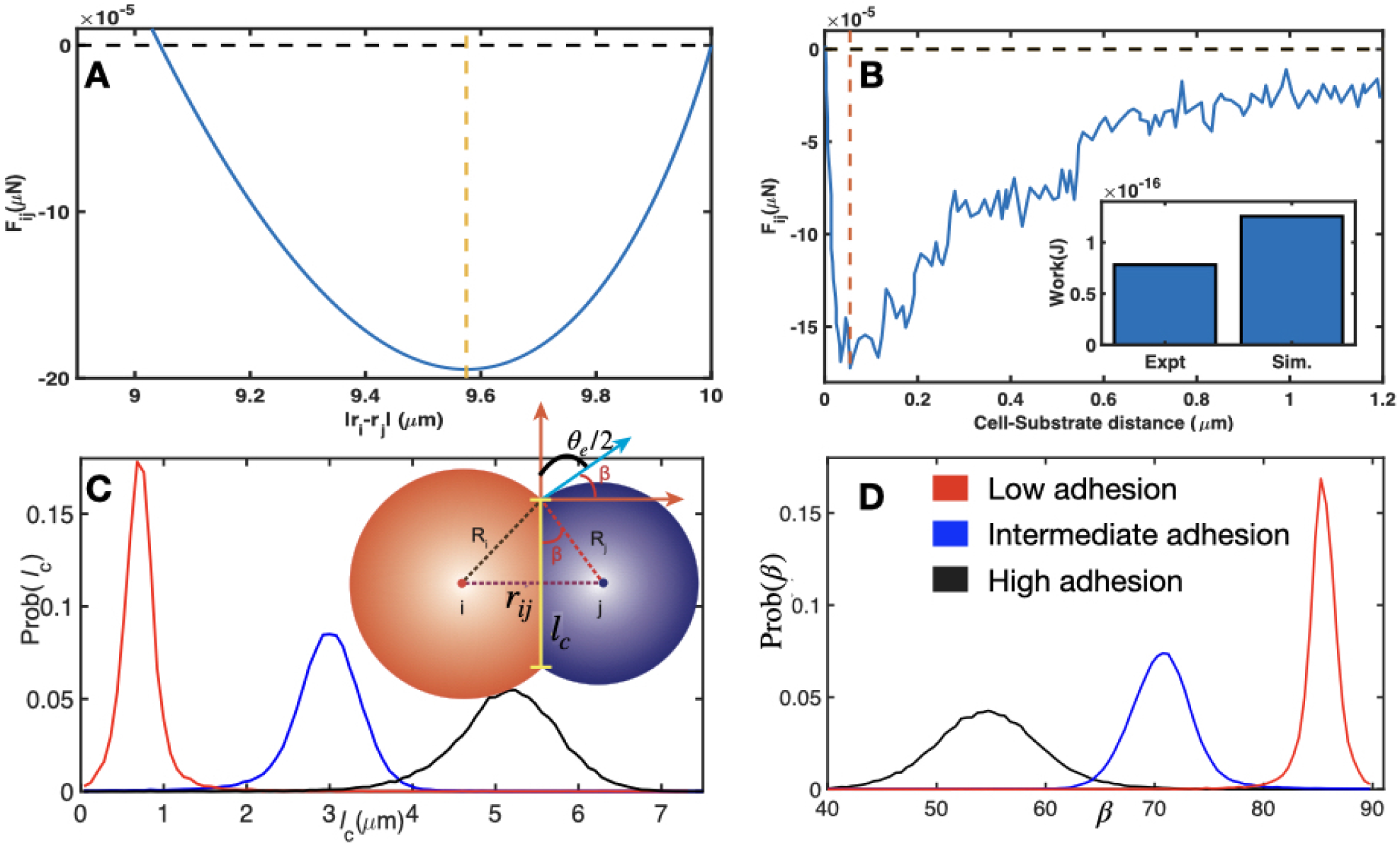
**A)** Force on cell *i* due to *j, F*_*ij*_, for *R*_*i*_ = *R*_*j*_ = 5 *µm* using mean values of elastic modulus, poisson ratio, receptor and ligand concentration (see Table I in the SI). *F*_*ij*_ is plotted as a function of cell center-to-center distance| **r**_*i*_− **r**_*j*_|. We used *f*^*ad*^ = 1.75 ×10^−4^*µ*N*/µ*m^2^ to generate *F*_*ij*_. **B)** Force-distance data extracted from single cell force spectroscopy (SCFS) experiment.^39^ Inset shows the work required to separate cell and E-cadherin functionalized substrate in SCFS experiment and two cells in theory, respectively. Minimum force values are indicated by vertical dashed lines. **(C)** Probability distribution of contact lengths between cells for *f*^*ad*^ = 0 (red), *f*^*ad*^ = 1.5 ×10^−4^ (blue) and *f*^*ad*^ = 3 ×10^−4^ (black). Inset shows how cell-cell adhesion dictates the angle of contact between cells, *β*, and the length of contact, *l* _*c*_. **(D)** Probability distribution of contact angles, *β* as a function *f*^*ad*^. The color scheme is the same as in **(C)**.

### Dependence of cell-cell contact length (*l*_*c*_) and contact angle (*β*) on *f*^*ad*^

Based on the center-to-center distance, *r*_*ij*_ = |*r*_*i*_ − *r*_*j*_|, between cells *i* and *j*, and the contact length, *l*_*c*_, the contact angle between two adhering cells, *β*, can be calculated (see Inset Fig.1**(C)** for the definition). Let *x* be the distance from center of cell *i* to contact zone marked by *l*_*c*_, along *r*_*ij*_. Similarly, we define *y* as the distance between center of cell *j* to *l*_*c*_ once again along *r*_*ij*_ (Fig. 1**(C)**). Based on the right triangle between *x, R*_*i*_ and 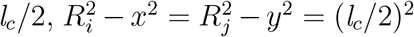 and *x* + *y* = *r*_*ij*_, we obtain,

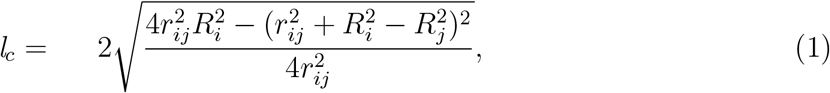

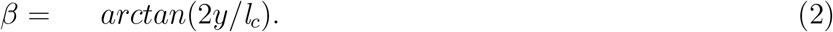

The probability distributions for *l*_*c*_ and *β* from the simulation as *f*^*ad*^ is varied are shown in Figs.1**(C)** and **(D)**. The extent of the overlap between cells increase as *f*^*ad*^ increases, leading to enhanced jamming at higher adhesion strengths. The distribution of *β* (see Fig.1**(D)**), whose width increases at higher adhesion strength, shows that the mean angle value decreases with *f*^*ad*^. By noting that 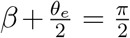 (*θ*_*e*_ is defined previously^40,41^) we infer from the results in Fig.1**(D)** that 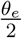 should increase as *f*^*ad*^ increases. Indeed, this is precisely what was found in phase transition in the zebrafish blastoderm (see Fig. 4C of Ref. ^41^ showing decreased connectivity with decreased cell-cell adhesion). The plots in Fig.1 show that the range of cell-cell adhesion strength we use in the simulations is in a reasonable range.

The dependence of the total cell-cell interaction force, *F*_*ij*_ as a function of *f*^*ad*^ is shown in Fig. 2**(A)**. Not unexpectedly, the location of the minimum in *F*_*ij*_ decreases upon increase in *f*^*ad*^ (see the inset in Fig. 2**(A)**). A schematic sketch of the enhanced adhesion with increasing *f*^*ad*^, due to greater E-cadherin expression, is shown in Fig. 2**(B)**.

**Figure 2:**
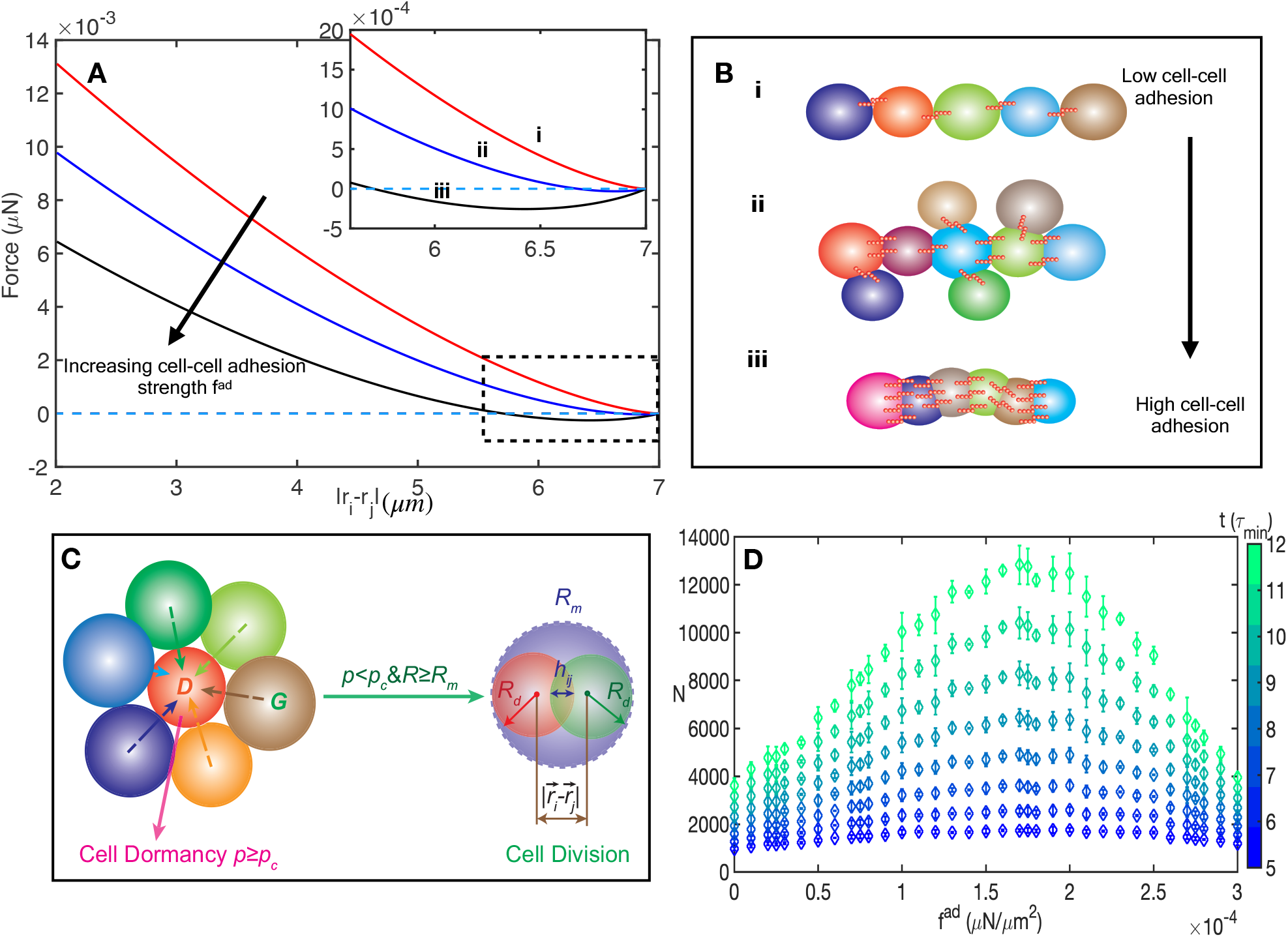
Forces and non-monotonic growth of the MCS. **(A)** The interaction force between cells as a function of cell center-to-center distance. As *f*^*ad*^ increases, the magnitude of the adhesive force increases. Inset shows the regime highlighted in the dashed box where *f*^*ad*^ increases from *i* (weakest) to *iii* (strongest). **(B)** Impact of three different regimes of *f*^*ad*^ on the spatial arrangement of the cells. Cadherin molecules are represented as short red bonds. The three adhesive regimes in the simulations correspond to (i) low, (ii) intermediate, and (iii) high values of *f*^*ad*^. **(C)** Cell dormancy (left) and cell division (right) in 3D cell collectives. If the local pressure *p*_*i*_ on the *i*^*th*^ cell (due to contacts with the neighboring cells) exceeds a the critical pressure *p*_*c*_, cell enters the dormant state, *D*. Otherwise, the cells undergo growth (*G*) until they reach the mitotic radius, *R*_*m*_ when the cell divides into two identical cells with identical radii *R*_*d*_. The value of *R*_*d*_ is determined to ensure that the total volume upon cell division is conserved. A cell that is dormant at a given time may transit from that state into growth phase at subsequent times. **(D)** Total number of cells (*N* in units of 1,000), from *t* = 5*τ*_*min*_ to 12*τ*_*min*_, at intervals of *τ*_*min*_(= 54, 000 sec), as a function of *f*^*ad*^) for *p*_*c*_ = 10^−4^MPa. The non-monotonic proliferation behavior is pronounced at *t* = 12*τ*_*min*_ (see Fig. 2**(D)**). Error bars here, and henceforth, are calculated from the standard deviation. The scale on the right gives time (*t*) in units of *τ*_*min*_.

### Physical and biological variables

The interaction parameters characterizing the systematic forces between cells are the two elastic constants (modulus - *E*_*i*_ and Poisson ratio - *ν*_*i*_), and *f*^*ad*^. In addition, cell birth (*k*_*b*_) and apoptotic rates (*k*_*a*_) are required to describe the evolution of the MCS. The values of *k*_*a*_ and *k*_*b*_ depend on the complicated biology governing cell fate, which are taken as parameters in the simulations. In the simulations, we chose the 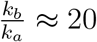 (see Table I in the SI) in order to model rapid tumor growth.

If *E*_*i*_, *ν*_*i*_, *k*_*a*_, and *k*_*b*_ are fixed, then tumor evolution is determined by *f*^*ad*^ and a specified threshold pressure, *p*_*c*_. If the pressure on the *i*^*th*^ cell at a given time (*p*_*i*_(*t*)) is less than *p*_*c*_ the cell grows, and eventually divides. If *p*_*i*_(*t*) *> p*_*c*_, the cell enters the dormant phase (Fig. 2**(C)**. We note parenthetically that *p*_*c*_ ≥ 0 is required for describing time dependent tumor growth law observed in experiments (see Fig. S1A in the SI). The existence of *p*_*c*_ above which cell division is halted is in agreement with recent experiment.^42^ Here, we explore the effects of *f*^*ad*^ and *p*_*c*_, on tumor proliferation.

## Results

### Cell proliferation depends non-monotonically on cell-cell adhesion strength

*f* ^*ad*^: We first quantify how changing *f*^*ad*^ impacts cell division, and hence the overall tumor growth. The simulations were initiated with 100 cells at *t* = 0, and the evolution of the cells is determined by the equations of motion, and the rules governing birth and apoptosis (see SI for a brief description). We performed simulations at different values of *f*^*ad*^ until ∼ 7.5 days (= 12*τ*_*min*_, where 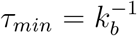 is the average cell division time). The number of cells (*N*) as a function of *f*^*ad*^, at various times in the range of 5*τ*_*min*_ to 12*τ*_*min*_ (with colors for time *t* in units of *τ*_*min*_), are shown in Fig. 2**(D)**. The central result, summarized in Fig. 2**(D)**, shows that tumor proliferation depends non-monotonically on *f*^*ad*^. The tumor growth is enhanced as *f*^*ad*^ increases from small values, attains a maximum at 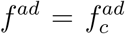, and decreases as *f*^*ad*^ exceeds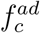.

On increasing *f*^*ad*^ from 0 to 1.75 × 10^−4^*µ*N*/µ*m^2^, the total number of cells, *N*, at *t* = 12*τ*_*min*_(∼7.5 days; green in Fig. 2**(D)**) increases substantially. At higher intercellular adhesion strengths (*f*^*ad*^ *>* 1.75 × 10^−4^), the proliferation capacity (PrC; quantified in terms of the total number of cells on day 7.5) is suppressed (Fig. 2**(D)**). While *N* = 12, 000 cells on day 7.5 at *f*^*ad*^ = 1.75 × 10^−4^*µ*N*/µ*m^2^, at higher *f*^*ad*^ values (such as 3 × 10^−4^*µ*N*/µ*m^2^), the tumor consists of only 4, 000 cells at the same time *t*. The surprising non-monotonic dependence of *N* on *f*^*ad*^, is qualitatively consistent with recent experiments,^39,43,44^ as we discuss below.

### Competition between local pressure and jamming drives non-monotonic proliferation

The non-monotonic tumor proliferation may be understood from the following physical arguments. Proliferation capacity is determined by the number of cells with pressure less than *p*_*c*_. If the pressure on a cell exceeds *p*_*c*_, it becomes dormant. Therefore, two factors should be considered in determining how mechanical feedback dictates cell proliferation: (i) dependence of intercellular pressure, and (ii) cell spatial packing, on *f*^*ad*^. Because the average pressure on a cell depends on the magnitude of the net adhesive force, the cell pressure decreases at higher values of *f*^*ad*^ (see SI Section I). This is similar to the pressure in real gases (van der Waals picture) in which the inter-particle attraction leads to a decrease in the average pressure. Therefore, within a certain range of 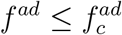, we expect the number of cells that is poised to divide should increase, causing enhanced tumor proliferation (van der Waals regime). At very high values of *f*^*ad*^, however, the adhesive interactions becomes so strong that the number of nearest neighbors of a given cell increases (Figs. S2**(A)** and S2**(B)** in the SI). This leads to an increase in pressure that decreases proliferation (jamming regime^6^). From these arguments it follows that cell proliferation would increase with increasing *f*^*ad*^ at low cell-cell adhesion strengths. However, at high *f*^*ad*^ cell proliferation would be limited due to a jamming effect.

The total average pressure (*p*_*t*_) that a cell experiences is given by 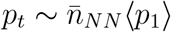 where 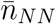 is the mean number of nearest neighbor cells, and ⟨*p*_1_⟩ is the average pressure a neighboring cell exerts on a cell. We use simulations to estimate the dependence of 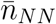 on *f*^*ad*^ (see SI Figs. S2**(A)**-**(B)**). For cell *i*, the nearest neighbors are defined as cells with non-zero overlap (*h*_*ij*_ *>* 0). To obtain ⟨*p*_1_⟩, we expand around *h*_0_, the cell-cell overlap value where both the attractive and repulsive interaction terms are equal 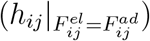, corresponding to the overlap distance at which *p* = 0 (see SI Fig. S1B). By expanding to first order, the average pressure of a single cell is 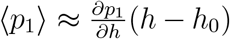, where, 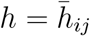, is the mean cell-cell overlap, dependent on *f*^*ad*^ (see SI Fig. S3). From the simulations, *h* − *h*_0_ increases approximately linearly with *f*^*ad*^ (see SI Fig. S4) implying that high *f*^*ad*^ limits cellular position rearrangements.

To investigate how cell spatial arrangement affects intercellular pressure, we calculate the pressure gradient with respect to cell-cell overlap, 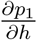, using

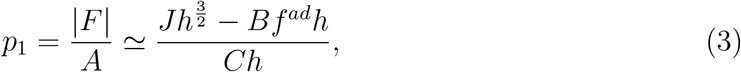

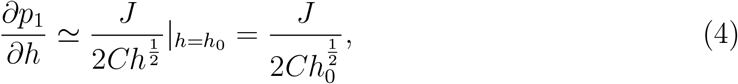

to separate out the dependence of pressure on *h*_*ij*_ and *f*^*ad*^. In Eqs. 3 and 4 *J, B* and *C* are independent of cell-cell overlap (*h*) and *f*^*ad*^. They are obtained from the definitions of elastic and adhesive forces and the definition of intercellular contact area, *A*_*ij*_. The resulting expressions are: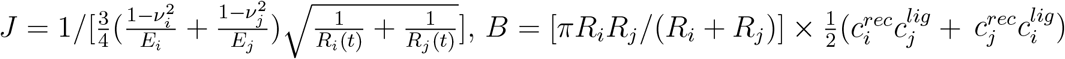, and *C* = *πR*_*i*_*R*_*j*_*/*(*R*_*i*_ + *R*_*j*_). On equating the repulsive and attractive interaction terms, we obtain *h*_0_ ≈ *K*(*f*^*ad*^)^2^, implying 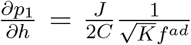, where, *K* = (*B/J*)^2^. Thus, the total pressure (*p*_*t*_) exerted on a cell is 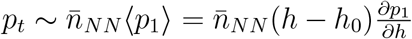, can be rewritten as,

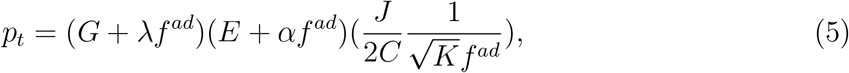

because the mean number of near-neighbors 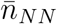 increases with *f*^*ad*^, which may be approximately written as *G* + *λf*^*ad*^ (see Fig. S2 A-B; *G* and *λ* are constants obtained from fitting simulation data). Similarly, the deviation of the cell-cell overlap from optimal packing (*h*−*h*_0_) increases linearly as *E* + *αf*^*ad*^ (see Fig. S4; *E, α* are constants). Notice that Eq. 5 can be written as, 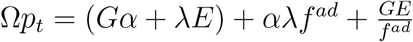, where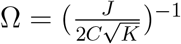. In this form, the second term depends linearly on *f*^*ad*^ and the third is inversely proportional to *f*^*ad*^. Since the enhancement in proliferation is maximized if the total pressure, *p*_*t*_, is minimized, the minimum in the total pressure on a cell is given by the solution to 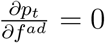. Therefore, the predicted optimal cell-cell adhesion strength is 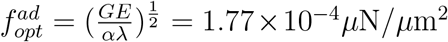. This is in excellent agreement with the simulation results for the peak in the tumor growth (*N* (*t* = 7.5 days) in Fig. 2**(D)**) at *f*^*ad*^ ≈ 1.75 × 10^−4^*µN/µm*^2^. More importantly, the arguments leading to Eq. 5 show that opposing contributions to the average pressure with increasing 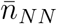 and decreasing intercellular pressure at high *f*^*ad*^ drives the non-monotonic proliferation.

### Pressure dependent mechanical feedback controls spatiotemporal proliferation patterns - Insights from simulations

We now consider how intercellular pressure varies with time and distance from the tumor center to periphery. Pressure experienced by a cell is highly dynamic, with peaks and subsequent drops in *p*_*i*_(*t*) (Figs. 3**A-C**). In Fig. 3**(A)**, black arrows highlight a switch in pressure from *p*_*i*_(*t*) *> p*_*c*_ (dormant phase) followed by *p*_*i*_(*t*) *< p*_*c*_ (growth phase). Such dynamic fluctuations in the individual cell pressure is independent of *f*^*ad*^ - as seen in Figs. 3A-C for *f*^*ad*^ = 0, *f*^*ad*^ = 1.5 × 10^−4^ and *f*^*ad*^ = 3 × 10^−4^ respectively. Notice that at early times 0 *< t*; ≲ 3.7*τ*_*min*_(∼ 200, 000*sec*), individual cell pressure values are lower on an average compared to later times *t ≲*7.4*τ*_*min*_(∼ 400, 000*s*). As the tumor cells continue to proliferate and become jammed, the pressure experienced by individual cells increase. At later times, the minimum individual pressure experienced by a cell tends to hover near the critical pressure as indicated by the dashed line in Figs. 3A-C.

**Figure 3:**
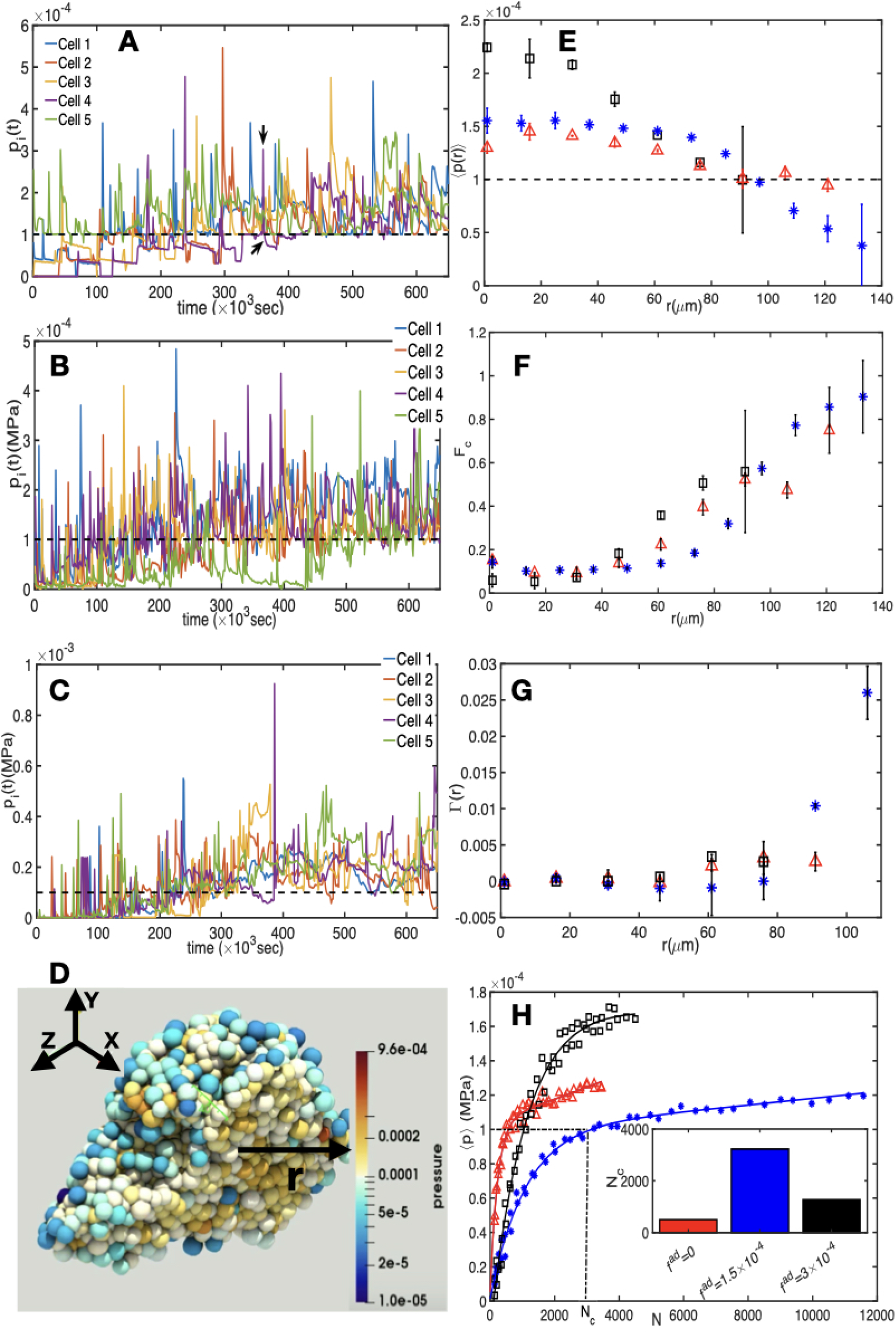
Temporal and spatial variations in pressure. **(A)**Pressure as a function of time for selected individual cells at *f*^*ad*^ = 0, **B)** *f*^*ad*^ = 1.5× 10^−4^ and **C)** *f*^*ad*^ = 3× 10^−4^. Dashed black lines correspond to the *p*_*c*_ value. Black arrows in **(A)** highlight *p*_*i*_ above *p*_*c*_. **(D)** A cross section through a tumor spheroid with distance *r* from tumor center to periphery at *t*∼ 12*τ*_*min*_ for *f*^*ad*^ = 0. Typically, pressure decreases from the core to the periphery of the MCS. **(E)** The average pressure experienced by cells at a distance *r* (*µ*m) from the MCS center for *f*^*ad*^ = 0 (red), *f*^*ad*^ = 1.5 ×10^−4^ (blue) and *f*^*ad*^ = 3 ×10^−4^ (black) (same symbols and color are used for panels **F-H**). The dashed line is *p*_*c*_ = 10^−4^MPa. Cells in the core are dormant with pressure above *p*_*c*_. **(F)** The fraction of cells with *p < p*_*c*_ as a function of *r*. At the tumor periphery cells are most likely not dormant, in agreement with **(E). (G)** Proliferation rate, Γ, is enhanced at the tumor periphery as seen in the increased values at distances *r* from the center of the MCS. The value of *t* = 12*τ*_*min*_ for **(E)**-**(G). (H)** Average pressure experienced by cells, ⟨*p ⟩*, as a function of the total number, *N*, of cells; ⟨ *p* ⟩ versus *N* at *f*^*ad*^ = 0 (red;triangle), *f*^*ad*^ = 1.5 × 10^−4^ (blue;asterisk) and *f*^*ad*^ = 3 × 10^−4^ (black;squares). The corresponding double exponential fits are, 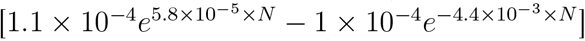 (red line, *f*^*ad*^ = 0), 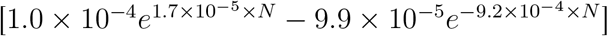 (blue line, *f*^*ad*^ = 1.5 ×10^−4^) and 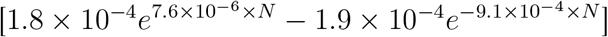 (black line, *f*^*ad*^ = 3 ×10^−4^). Inset shows *N*_*c*_, the total number of cells at which the average pressure evolves to the critical pressure ((*p*) → *p*_*c*_).

Next, we quantified the experimentally measurable^45,46^ spatial patterns in pressure experienced by cells within the MCS. The pressure associated with cells at the MCS center exceeds *p*_*c*_ (see Fig. 3**(D)**) leading to suppressed cell growth and division. In contrast, as visualized in Fig. 3**(D)** and quantified in Fig. 3**(E)**, the average pressure experienced by cells close to the tumor periphery tend to be below *p*_*c*_. The *f*^*ad*^ dependent average pressure at *r* (distance from tumor center) is quantified using,

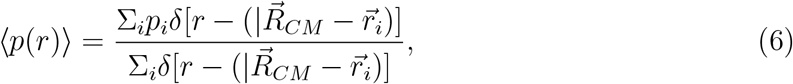

showing that the lowest average pressure at the tumor periphery is at the intermediate value of *f*^*ad*^ = 1.5 × 10^−4^ (see Fig. 3**(E)**). Due to the high pressure experienced by cells near the tumor center, we expect a well defined spatial trend in the fraction of proliferating cells, *F*_*c*_ = *N* (*p*_*i*_ *< p*_*c*_)*/N*, where *N* (*p*_*i*_ *< p*_*c*_) is the number of cells with pressure *p*_*i*_ *< p*_*c*_. Indeed, the proliferating cell fraction is low (*F*_*c*_ *<* 0.2) at the MCS center compared to the periphery where it approaches unity (*F*_*c*_ → 1), indicating that cells are in the growth/division phase (see Fig. 3**(F)**).

There is a rapid increase in *F*_*c*_ near the tumor periphery for *f*^*ad*^ = 1.5 × 10^−4^ (see the blue asterisks in Fig. 3**(F)**). To better understand the spatial dependence on proliferation, we calculated the average cell proliferation rate, 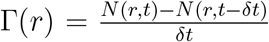, where *N* (*r, t*) is the number of cells at time *t* ≈ 12*τ*_*min*_ and *δt* ≈ 0.1*τ*_*min*_ is the time interval, as a function of distance *r* from the tumor center. The average is over the polar and azimuthal angles, for all the cells between *r* and *r* + *δr*. Closer to the tumor center, Γ(*r*) ∼ 0, indicating no proliferative activity. For larger values of *r*, approaching the MCS periphery, the proliferation rate increases rapidly. The cell proliferation rate is similar for different *f*^*ad*^ at small *r* (tumor core), while a much higher proliferation rate is observed for the intermediate value of *f*^*ad*^ at large *r* (see Fig. 3**(G)**). The simulated spatial proliferation profile is in agreement with experimental results,^45^ where increased mechanical stress is correlated with lack of proliferation within the MCS, albeit when stress is exerted externally. We note that even in the absence of an external applied stress, intercellular interactions can give rise to heterogeneity in the spatial distribution of intercellular mechanical stresses and impact spatial proliferation patterns.

Our results show that the non-monotonic cell proliferation behavior emerges because the fraction of cells with pressure less than *p*_*c*_ is localized at the MCS periphery. The average pressure, ⟨*p* ⟩, experienced by cells as a function of *N*, at the three values of *f*^*ad*^, is shown in Fig 3**(H)**. *N*_*c*_, defined as the number of cells at which ⟨*p* ⟩ = *p*_*c*_, exhibits a nonmonotonic behavior (inset in Fig 3**(H)**), which supports the maximum number of cells at an intermediate *f*^*ad*^. We surmise that, at intermediate *f*^*ad*^, cells spatially rearrange and pack effectively to minimize the average cell pressure. For all three *f*^*ad*^ values, an initial regime where pressure rises rapidly is followed by a more gradual increase in pressure, coinciding with the exponential to power law crossover in the growth in the number of cells (see Fig. S1A). The average pressures (⟨*p* ⟩) as a function of *N* is well fit by double exponential functions (Fig 3**(H)**). Hence, by experimentally measuring the cell proliferation rate, we can delineate the pressure sensing mechanism of cells.

### Fraction of cells with pressure less than *p*_*c*_ controls non-monotonic proliferation

The pressure experienced by a cell as a function of contact with other adhering cells plays an essential role in determining the observed non-monotonic proliferation. For *f*^*ad*^ *>* 0, there is a minimum in the pressure at a non-zero cell-cell overlap distance, *h*_*ij*_ (SI Fig. S1B). For instance, at *f*^*ad*^ = 1.5 × 10^−4^*µ*N*/µ*m^2^ and 3 × 10^−4^*µ*N*/µ*m^2^, the mean pressure exerted on the cells is zero at *h*_*ij*_ ∼ 0.4*µ*m and 1.3*µ*m respectively. The minimum pressure is a consequence of the balance between adhesive and elastic forces. At the minimum pressure (*p*_*i*_ → 0), the proliferation capacity (PrC) of the cells is maximized because the cells are almost always in the growth phase. However, the impact of variations in individual cell pressure on growth of the cell collective is unclear. To determine the effect of *f*^*ad*^ on proliferation we first quantify the spatial and time dependent evolution of pressure.

The fraction of cells, *F*_*c*_ in the proliferating regime, corresponding to the shaded portion in Fig. 4**(A)** exhibits a non-monotonic behavior (inset in Fig. 4**(A)**) with a peak at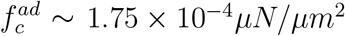. The *F*_*c*_ value in the growth phase (*p*_*i*_ *< p*_*c*_) peaks at ≈ 38%, between 1 × 10^−4^ *µ*N*/µ*m^2^ *< f*^*ad*^ *<* 2 × 10^−4^ *µ*N*/µ*m^2^. At lower and higher values of *f*^*ad*^, *F*_*c*_ is at or below 25% on day 7.5. The inset in Fig. 4**(A)** makes it clear that *F*_*c*_, which determines the proliferative capacity of the MCS, depends non-monotonically on *f*^*ad*^.

**Figure 4:**
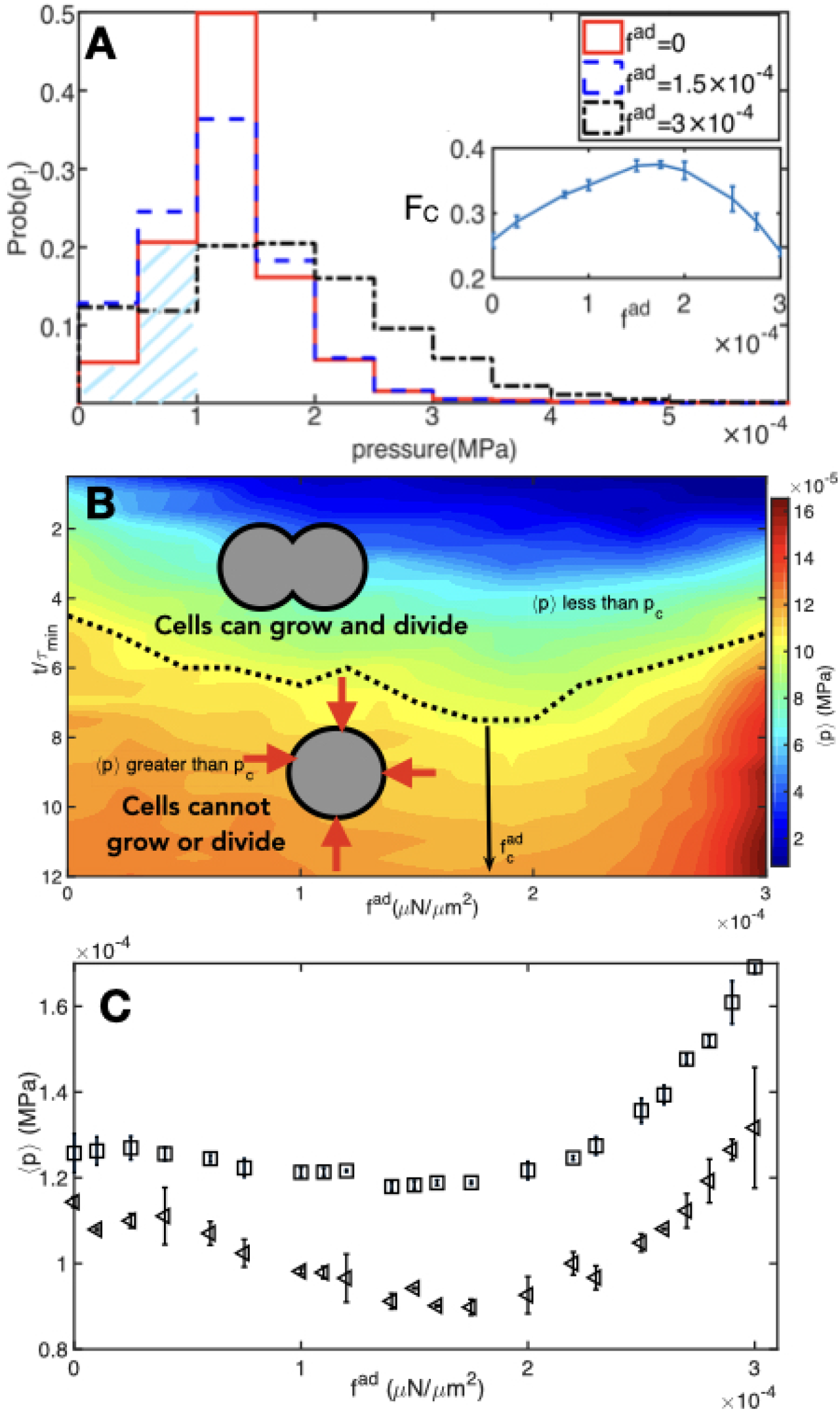
**(A))** The probability distribution of pressure exerted on the cell on day 7.5 for three different values of *f*^*ad*^. The shaded regions represent cells with pressure below *p*_*c*_ = 1 ×10^−4^MPa. Fraction of cells (*F*_*C*_), with *p < p*_*c*_, that grows and proliferate is shown in the inset. **(B)** Phase plot of the average pressure on the cells as a function of time and *f*^*ad*^. The pressure scale is on the right. The dashed line is the boundary between the regime when cells on an average grow and divide versus the regime where mechanical pressure restricts cell growth. **(C)** The average pressure ⟨ *p* ⟩ experienced by cells at *t* = 6*τ*_*min*_ (left triangles; day ∼3.75 of growth) and *t* = 12*τ*_*min*_ (squares; ∼day 7.5 of growth) for different values of *f*^*ad*^.

The results in Fig. 2**(D)** and the inset in Fig. 4**(A)** suggest a phase diagram, shown in Fig. 4**(B)**, in terms of the average cell pressure 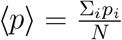, color map) and *f*^*ad*^. The dotted line is the boundary, ⟨ *p* ⟩ = *p*_*c*_, between the regimes where cells on an average grow and divide (*p*_*i*_ *< p*_*c*_), and where they are dormant (*p*_*i*_ *> p*_*c*_). At a fixed *t*, the regime between 1 × 10^−4^ ≤ *f*^*ad*^ ≤ 2 × 10^−4^ shows a marked dip in the average pressure experienced by the cells. The (*p*) *< p*_*c*_ regime becomes pronounced between 2*τ*_*min*_ ≤ *t* ≤ 7*τ*_*min*_. In Fig. 4**(C)**, the average pressure, ⟨ *p* ⟩, at *t* = 6*τ*_*min*_ and 12*τ*_*min*_ as a function of *f*^*ad*^, has a minimum between *f*^*ad*^ ≈ 1.5 − 2 × 10^−4^ *µ*N*/µ*m^2^. Clearly, there is a close relationship between proliferation and the local pressure on a cell. As the number of cells with pressure below *p*_*c*_ increases, the proliferation at cell collective level is enhanced. The origin of the minimum in the average pressure at 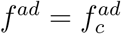 (Figs. 4**(B)** and **(C)**) is less clear. The theoretical explanation for this key finding was provided above based on Eq.5.

## Discussion

### Intercellular pressure determines the biomechanical feedback

Our simulations show that non-monotonic proliferation is determined by the fraction of proliferating cells, *F*_*C*_, with pressure less than *p*_*c*_. This picture is reminiscent of the mechanical feedback as a control mechanism for tissue growth proposed by Shraiman^11,17^ but with a key difference. In earlier work, cellular rearrangements does not occur readily, and the tissue is treated as an elastic sheet that resists shear^11^ with mechanical stresses serving as a feedback mechanism to restrict proliferation. In our case, large scale cell rearrangements are possible^34^ as a cell or group of cells could go in and out of dormancy. Despite the differences, the idea that local pressure serves as a regulatory mechanism of growth could be a general feature of mechanical feedback.^11^

### Comparison with experiments

We now analyze some experiments in light of predictions from our simulations. Based on the observation of enhanced cell migration at low E-cadherin levels, tumor invasion and metastasis is considered to be a consequence of loss of E-cadherin expression.^47^ However, most breast cancer primary and metastatic tumors express E-cadherin.^48^ Our simulations suggest that tumor spheroids made up of cells with higher E-cadherin expression (E-cad(control)) grow ≈ 1.9 times faster compared to tumor spheroids made up of low E-cadherin (E-cad(-)) expressing cells (see Fig. 5**(A)**), in reasonable agreement with experiments.^44^ Comparison of the tumor diameter growth between E-cad(-) cells and E-cad(control) (red and blue lines respectively, Fig. 5**(A)**) shows enhanced growth rates for E-cad(control) cells at *t >* 4 days. The linear fit for the simulated tumor diameter growth rate is obtained by analyzing data at *t >* 4 days (dashed lines; Fig. 5**(A)**). The growth rate ratio is then calculated by taking the ratio of the slope of the two diameter growth lines.

**Figure 5:**
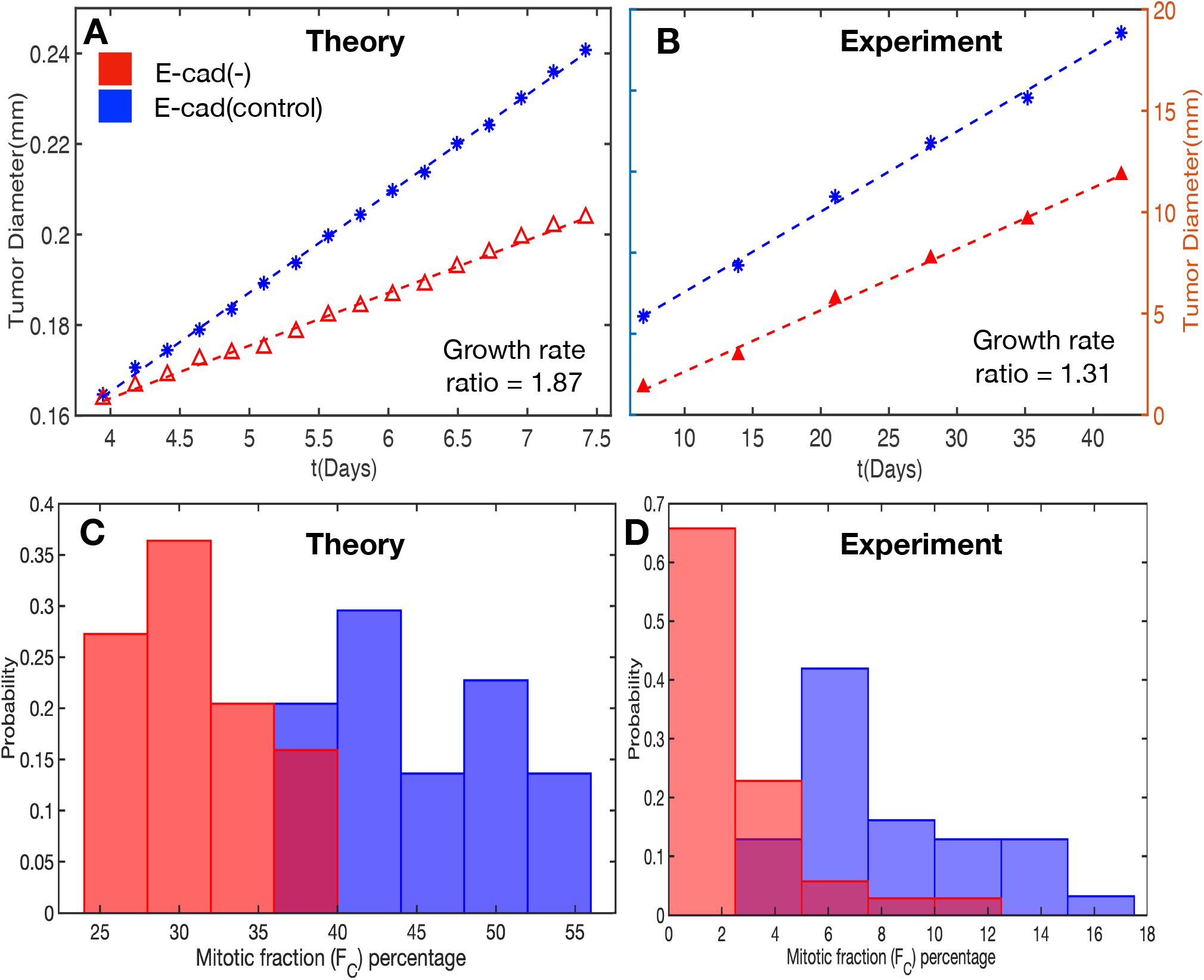
Comparison between simulations and experiments. **(A)** Kinetics of growth in the diameter of the MCS composed of cells with low E-cadherin expression (red; *f*^*ad*^ = 0; E-cad(-)) and intermediate E-cadherin expression (blue; *f*^*ad*^ = 1.75× 10^−4^; E-cad(control)) from simulations show enhanced tumor growth rate due to higher E-cadherin expression. The tumor diameter growth is linear at *t >* 4 days, independent of the E-cadherin expression level. However, the growth rate of the tumor colony at intermediate E-cadherin expression is larger in agreement with experimental results. **(B)** Growth rates of the longest tumor dimension in low E-cadherin expressing organoids compared to control organoids with normal E-cadherin expression from experimental data (data was extracted from Ref.^44^). **(C)** Calculated probability distribution of the mitotic fraction, *F*_*C*_, in cell colonies expressed as % for low E-cadherin tumor MCS (red) and normal E-cadherin expressing tumor MCS (blue) shows enhanced proliferation capacity when E-cadherin expression is increased. **(D)** In agreement with the theoretical predictions, the probability distribution of the mitotic fraction, *F*_*C*_ for low E-cadherin tumor organoids (red) and normal E-cadherin expressing tumor organoids (blue) shows enhanced proliferation capacity when E-cadherin expression is increased. Data was extracted from experiments (see the SI data in Table 3e in the experimental report Ref.^44^). Although there is paucity of experimental data, it is encouraging that the currently available measurements are in line with our predictions.

Padmanaban et. al^44^ compared the tumor growth behavior between E-cadherin expressing cells (control) and E-cadherin negative cells (characterized by reduced E-cadherin expression compared to control) using 3D tumor organoids. They conclude that tumors arising from low E-cadherin expressing cells (E-cad(-)) were smaller than tumors from control E-cadherin (E-cad(control)) expressing cells at corresponding time points over multiple weeks of tumor growth (see Fig. 5**(B)**). By extracting the tumor growth rate for E-cad(-) and E-cad(control) organoids from the experimental data,^44^ the longest tumor dimension for E-cad(control) tumor organoids expanded ≈ 1.3 times faster compared to E-cad(-) tumor organoids (see Fig. 5**(B)**). Both the linear functional form of the tumor growth rates and the higher growth rates in E-cad(control) tumor spheroids as compared to E-cad(-) tumor spheroids are in reasonable agreement with the simulations. It is worth emphasizing that the simulation results were obtained without any fits to the experiments.^44^

To ensure that the difference in tumor growth rate is due to differences in cell proliferation, we analyzed the ratio of the number of actively dividing cells to the total number of cells with experimental data obtained from staining tumor cell colonies for PH3 (phospho-histone 3; a mitotic marker). As predicted, E-cadherin expressing tumor cell colonies have a larger mitotic fraction (blue histogram, Fig. 5**(C)**) compared to low E-cadherin expressing tumor cells (red histogram, Fig. 5**(C)**). We compare the mitotic fraction, *F*_*C*_, in E-cad(-) and E-cad(control) in the Figs. 5**(C)**-**(D)** between theory and experiment. In agreement with our simulation predictions, enhanced mitotic fraction is seen in tumor organoids with normal E-cad expression^44^ (see Fig. 5**(D)**).

## Conclusions

We have established that increasing cell-cell adhesion strength contributes to mechanical pressure based inhibition in cell growth and proliferation. Surprisingly, growth of the multicellular spheroid exhibits a non-monotonic behavior as a function of increasing cell adhesion strength. Cell-cell adhesion strength and critical pressure based inhibition of cell growth, qualitatively captures cell proliferation patterns observed in the context of cancer progression.

The observed role of E-cadherin in tumor growth could be related to a feedback mechanism due to changes in local pressure as the cell-cell interaction strength is varied. The pressure feedback on the growth of cells accounts for cell proliferation in the simulations.

In actuality, it may well be that mechanical forces do not directly translate into proliferative effects. Rather, cell-cell contact (experimentally measurable through the contact length *l*_*c*_, for example) could biochemically regulate cell signaling (eg: Rac1/RhoA), which in turn controls proliferation, as observed in biphasic proliferation of cell collectives in both two and three dimensions. ^49,50^ Nevertheless, contact inhibition of proliferation, based on mechanical pressure experienced by single cells can be captured using a pressure based feedback on proliferation in computer simulations. Such mechanical control of proliferation could be a generic mechanism governing how cell-cell adhesion strength, acting on the scale of at best few microns, affects tumor growth that occurs on the scale of millimeters.

## Acknowledgements

We acknowledge Anne D. Bowen at the Visualization Laboratory (Vislab), Texas Advanced Computing Center, for help with video visualizations. We are grateful to Dr. Paul Langridge for useful comments. This work is supported by the National Science Foundation (PHY 17-08128) and the Collie-Welch Chair through the Welch Foundation (F-0019). A.N.M-K acknowledges funding support from the College of Science and Mathematics at Augusta University. The authors declare that they have no conflict of interest.

